# Effects of the nemertean egg predator *Carcinonemertes carcinophila* (Carcinonemertidae: Monostilifera) on the reproductive output and fecundity of the Florida Atlantic blue crab *Callinectes sapidus*

**DOI:** 10.64898/2026.10.19.760229

**Authors:** Max Taylor, Lucas Jennings, Donald C. Behringer

## Abstract

The blue crab, *Callinectes sapidus*, is an important crab fishery in Florida, valued at approximately $14.5 M in 2023. However, female blue crabs are host to the nemertean worm *Carcinonemertes carcinophila,* which preys upon its eggs. Other nemerteans from this genus have been linked to the collapse of fisheries, such as the Dungeness crab fishery in the 1960s and 1970s and the red king crab fishery in the 1980s. In 2022, the newly-discovered nemertean *Carcinonemertes conanobrieni* was found to parasitize the egg masses of female Caribbean spiny lobsters, *Panulirus argus,* and has had drastic impacts on the reproductive output and fecundity of this species. We hypothesized that nemertean egg parasites might have similar impacts on blue crabs. To investigate this, we collected 52 ovigerous female blue crabs from Jacksonville and Ormond Beach, 61.5% of which were infested with *C. carcinophila*. Infestation status did not significantly affect fecundity or reproductive output; instead, body mass and embryo stage were the strongest predictors. Infestation was more prevalent in crabs with late-stage embryos, consistent with nemertean life cycle timing. While most hosts carried few worms, a subset showed high infestation intensity, and both infestation status and nemertean density were significant predictors of embryo loss. This is a positive sign for the Florida blue crab fishery, as it suggests that the blue crab nemertean might not be as impactful to that fishery as others.

## INTRODUCTION

The Atlantic blue crab is an important Florida fishery, both commercially and recreationally (Steele 1998). In 2024, the commercial Florida blue crab fishery was valued at just over $16 M (USD) (FWC 2023). Beyond Florida, *C. sapidus* supports robust fisheries in Louisiana, Maryland, Virginia and North Carolina (Hill et al. 1989; Guillory & Elliot 2001; Walters et al. 2025). Ecologically, blue crabs are an integral part of benthic communities. As both predator and prey, blue crabs play a key role in ecosystem composition. Through its foraging behaviors, blue crabs have the ability to disrupt benthic sediments, affecting community organization and distribution of benthic prey species (Virnstein 1977; Martin et. al 1989; Hines et. al 1990; Perry et al. 2022). This species has a large disjunct distribution that ranges from Maine to Argentina (Santos & D’Incao 2004; Rodrigues et al. 2017), where it holds vital positions in Atlantic commercial fisheries and ecological communities.

Upholding fishery health and economic profitability starts with assessing the fertility of female crabs. Blue crab fishermen in Florida are required to release any ovigerous females they trap because they bear the fertility burden (FWC 2023), and it is highly encouraged that they also release female crabs without eggs as well because of the importance of females in maintaining the population. Female crab life history includes a terminal molt, during which she mates only once in their life (Darnell et al. 2009). The female stores the sperm she receives in her spermatheca (seminal receptacles), where it is later used to fertilize her embryos. Although male crabs are capable of mating multiple times per season, their population is vulnerable to reproductive stressors that affect reproduction. Studies suggest that blue crab reproduction can be limited by sperm quality (Hines et al. 2003) and quantity (Jivoff 2003). The most crucial detriment to decapod crustacean reproduction, however, starts with oviposition (Kuris 1990), a critical moment in a female’s reproductive cycle that marks the transition from the relative safety of an unfertilized egg to a fertilized embryo exposed to environmental dangers.

Egg predation is one of the leading causes of decapod crustacean fishery degradation (Kuris, 2020) and nemertean worms from the family Carcinonemertidae are among the most voracious (Kuris and Wickham, 1987). Their intense predation on crustacean brood masses has been implicated in the collapse of several fisheries, including the Dungeness crab in the 1960s and the red king crab in the 1980s (Wickham, 1979, Shields et al., 1989). The egg predator *Carcinonemertes regicides* was correlated with an average of 90% egg mortality in red king crabs in regions with outbreaks (Kuris et al. 1991). Nemertean worms have recently been discovered in other species, including the spiny lobster, *Panlinus argus* in 2017 (Simpson et al. 2017) and the Chilean lemon crab, *Cancer porteri* and a rock crab, *Romaleon setosum* in 2021 (Leiva et al. 2021). All three of these species are commercially important and decreases in fecundity could have severe economic consequences. In 2022,infestation in spiny lobsters by the egg predator *C. conanobrieni* resulted in a significant decline in fecundity and reproductive output (Berben et al. 2023; Baeza et al. 2026). This coupled with its 100% prevalence in Florida populations could lead to heavy impacts on the spiny lobster fishery (Stephens et al. 2025). Novel species of nemerteans have also been discovered in areas without prior infestations, such as the newly described *C. camanchaco*, which was found in two crab species in the southeastern Pacific where previously no nemerteans had been recorded (Leiva et al. 2021).

There are 22 species of Carcinonemertidae that have been recorded to feed on the embryos of over 60 crustacean species and their host specificity varies among species (Segonzac and Shields 2007). The Atlantic blue crab *Callinectes sapidus* is infected by the generalist nemertean worm *C. carcinophila,* which parasitizes 28 species of decapod crustaceans (Segonzac and Shields 2007); however, many of these findings lack genetic confirmation. Carcinonemertids have a short proboscis with a small stylet that they use to pierce the embryo and ingest the yolk, changing the color of the worm from cream to red-orange (Messick et al. 1992, McDermott et al. 1985). After mature *C. carcinophila* mate and lay their eggs in the crab brood mass, they migrate to the host’s gills (Schneider et al. 2023, Overstreet 1978). The gills provide a refuge for mature worms until the crab oviposits her next batch of embryos. Worm larvae either remain in the brood mass to prey on *C. sapidus* embryos or are expelled into the water column where they presumably may infest a new ovigerous female (Kuris 1993, Davis 1965).

Despite the commonality of *C. carcinophila* in blue crabs, the impact on their hosts’ fecundity has only recently been considered in the Chesapeake Bay. In Florida, the impact of carcinonemertid infestation on blue crab fertility has never been assessed. The lower latitudes and warmer water allow Florida blue crabs to remain active and reproduce nearly year-round (Steele 1998). Florida blue crab reproduction follows its own patterns and cycles, distinct from those in the Chesapeake Bay region, meriting separate reproductive surveys.

Previous studies have estimated the average fecundity of female blue crabs on Florida’s east coast at 2.1 - 2.2 million embryos per season (Hart et al. 2021). Blue crabs are multiparous, producing multiple broods per season. In Florida, primary spawning seasons occur during the late spring and late summer (Steele and Bert 1994). Florida blue crabs are estimated to produce between 6-7 broods per year and 18 broods per lifetime, while those in Chesapeake Bay produce 1-3 broods per year and 6-7 broods per lifetime (Hines et al. 2003). In contrast to Florida blue crabs, Chesapeake Bay blue crabs produce only around 900,000 embryos per brood (Hines et al. 2003).

Reproductive output differs greatly with climate, affecting mating seasons and activity levels. Given Florida’s extended reproductive season, a nemertean egg predator such as *C. carcinophila* could therefore possibly impact Florida blue crabs more than those in the Chesapeake Bay. Despite this potential threat, the impact of *C. carcinophila* on the fecundity and reproductive output of Florida blue crabs remains unexamined. To address this gap, we conducted an analysis to assess infestation extent. We used a subsampling technique and quantified the intensity of infestation, prevalence and embryo mortality, providing the first evaluation of *C. carcinophila’s* impact on Florida’s blue crab fishery.

## MATERIALS AND METHODS

### Sample Collection

Ovigerous *Callinectes sapidus* females were collected between January and February 2023 on Florida’s Atlantic coast from two locations; the St. Johns River (30.43043°N, 81.45544°W) and the Halifax River (N 29°12.047’ W 81°00.456’). Crabs were trapped with commercial blue crab traps by local fishermen. Crabs were then transported live to the Aquatic Pathology Laboratory at the University of Florida (Gainesville, Florida, USA). In the laboratory, crabs were anesthetized in an ice bath before processing. To obtain reproductive measurements, live crabs first had their pleopods detached and placed into Whirlpaks filled with natural seawater and frozen at -20°C. Some crabs autotomized legs during transportation, so we removed all limbs from all crabs prior to weighing. The crab bodies were then dried in a drying oven at 72°C for 72 hours.

### Fertility in Callinectes sapidus

To assess fertility in *C. sapidus,* we determined fecundity and reproductive output via a subsampling method adapted from Baeza et al. 2016. Embryo masses were allowed to thaw in cold water prior to subsampling. Under a Leica S8APO dissecting microscope, approximately 1000 embryos were haphazardly selected from four pleopods from an individual female. These embryos were counted as either viable, dead, or eaten. Dead embryos displayed a noticeable opaque hue different from their peers, and eaten embryos appeared to have no contents left, with the remaining capsule contorted into a collapsed football shape. Embryo masses were classified into four different stages microscopically in accordance with Hines et al. (2003); early no-eyed embryos, late no-eyed embryos, early eyed embryos, and late eyed embryos. These were later divided more simply into early stage, or no-eyed embryos, and late stage, or eyed embryos. Each of the subsamples was then dried in a drying oven at 72℃, along with the rest of the brood mass. After at least 72 hours, dried embryo subsamples and brood masses were weighed using an analytical balance.

Fecundity was defined as the total amount of viable embryos present in an entire brood mass. To calculate the total amount of embryos, both viable and dead, we used the following equation:

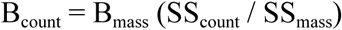

Where B_count_ = the number of viable and dead embryos in the brood mass, B_mass_ = the mass of the dried brood mass, SS_count_ = the number of viable and dead embryos in the four subsamples, and SS_mass_ = the mass of the four subsamples. Since eaten eggs contained no yolk and were present in small numbers, we concluded that they would have no significant impact on the brood mass. The ratio of viable to total embryos was applied to the B_count_ number to determine the total number of viable embryos in the brood mass, or the fecundity, with the following equation:

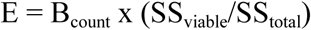

Where E = fecundity, SS_viable_ = the count of viable embryos across all four subsamples, and SS_total_ = the count of viable and dead embryos across all four subsamples. The number of embryos in the subsamples was also included in total fecundity count. We used a linear regression to determine the relationship between carapace width, crab body mass, and embryo stage with fecundity. In transporting crabs from the field to the lab, some of the lateral carapace spines broke off, which would lower measured carapace width. To account for this, we used crab mass to approximate crab size.

Reproductive output is defined as the ratio of embryo mass to the total crab body mass. We used a general linearized model to determine the relationship between reproductive output, embryo stage and infestation status.

### Species confirmation

Prior to the impact assessment After being removed from the egg mass, carcinonemertids were initially identified based on morphological traits. A DNA extraction was then performed on a whole, single adult using the Qiagen DNeasy Blood and Tissue kit (Hilden, Germany) according to the manufacturer’s to manufacturer’s protocol for animal tissue. The resulting extract was shipped to Novogene, California on dry ice for sequencing. A NEBNext Ultra DNA Library Prep Kit was used for library preparation prior to the sample being loaded on a NovaSeq 6000 (Illumina). Paired end metagenomic sequencing was performed using the 150 bp NovaSeq 600 SP reagent Kit (300 cycles) and the resulting sequencing dataset was sent to the University of Florida for bioinformatic analysis. The data was trimmed and quality checked using Trimmomatic v.0.36 (Bolger 2014), set to the default parameters. SPAdes v.4.0.0 (Bankevich et al. 2012) was used to assemble the paired and unpaired reads with default parameters and k-mer lengths: 21, 35, 55, 77 and 99. Metaxa2 v.2.2.3 (Bengtsson-Palme et al. 2015) was then used to identify the mitochondrial genome of the sample which was uploaded to CLC genomics workbench v.12 for circularization. The circularized mitogenome was then uploaded to the EU Galaxy webserver and MITOS2 was used to identify the *cytochrome oxidase 1* (COI) sequence. The resulting *cox1* sequence was then blasted using NCBI BLASTn. Sequences for 21 carcinonemertid species were downloaded from the NCBI database and MAFTT aligned (Katoh and Standley, 2013). Two species from the nemertean genus *Tubulanus* were used as an outgroup. A maximum likelihood analysis was carried out on the IQ-tree webserver (Trifinopoulos, 2016) where the most appropriate evolutionary model was chosen based on Bayesian inference criterion (BIC) (TIM3+F+I+G4). A maximum likelihood tree was computed using 1,000 bootstrap replicates over 1,536 nucleotide positions with a log score of -5664.02. The resulting tree was annotated using FigTree v1.4.4

### Presence of C. carcinophila in C. sapidus

To assess the impact of nemertean worms on *C. sapidus* in Florida, we inspected brood masses for the presence and density of worms, and worm impact. Positive infestation status was determined by presence of worms, mucus tubes, or eaten embryos. During fertility measurements, the embryo subsamples were also inspected for adult and juvenile worms. The density of *C. carcinophila* was measured as the number of worms per 1000 embryos. Embryo mortality was defined as the number of eaten embryos per 1000 embryos. Worm impact was calculated as a ratio between the number of worms and the embryo mortality. A linear regression was used to determine the relationship between embryo stage and infestation status. General linearized models were used to determine the relationship between nemertean density, number of eaten embryos and embryo stage.

## RESULTS

Out of the 52 crabs surveyed, 32 were infected with *C. carcinophila*, resulting in a 61.5% prevalence over all sites surveyed. At the St. John’s River location, 11 out of 20 crabs showed signs of infestation, or a 55% prevalence. At the Halifax River location, 21 out of 32 crabs showed signs of infestation, or a 66% prevalence. After initial morphological identification, all carcinonemertid species were confirmed as *Carcinonemertes carcinophila* after DNA extraction and *cytochrome oxidase 1* (COI) BLAST (Fig. 1). The mean fecundity for all locations was 1.75 ± 0.66 M embryos, and 1.85 ± 0.56 M at the Halifax River and 1.6 ± 0.74 M at the St. Johns River. The most significant predictor of fecundity was crab body mass (Fig. 2A, Table 1), but carapace width was also significant (Fig. 2B, Table 1). Infestation status was not a significant predictor of fecundity, even when embryo stage and crab body mass were added as covariates (Fig. 3, Table 1).

**Figure 1.**
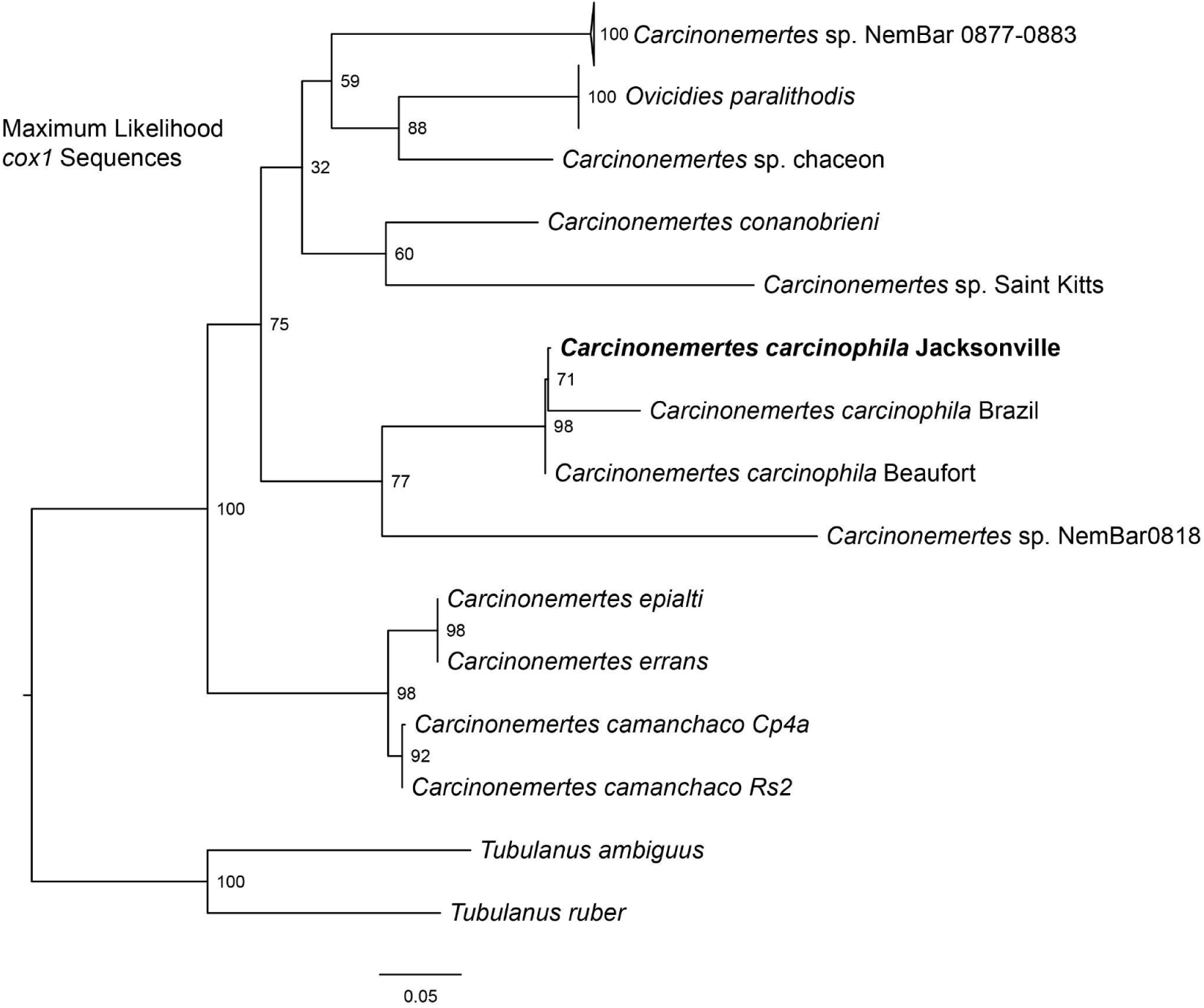
A maximum likelihood tree using the cytochrome oxidase subunit 1 (*cox1*) gene.

**Figure 2.**
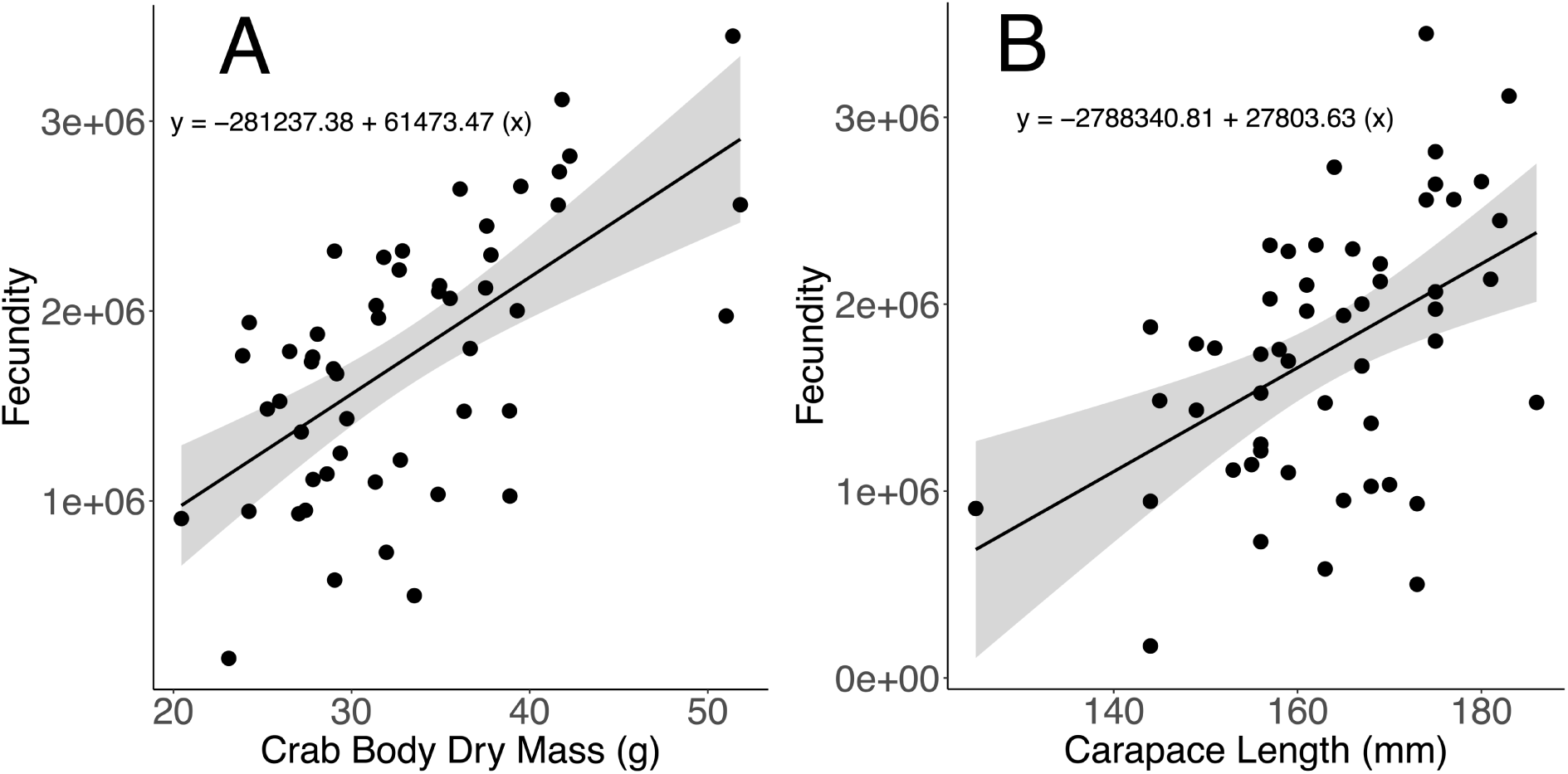
Effect of body mass on fecundity (A). Linear regression, R^2 = 0.338, p-value <0.01. Effect of carapace width on fecundity (B). Linear regression, R^2 = 0.32, p-value <0.01.

**Figure 3.**
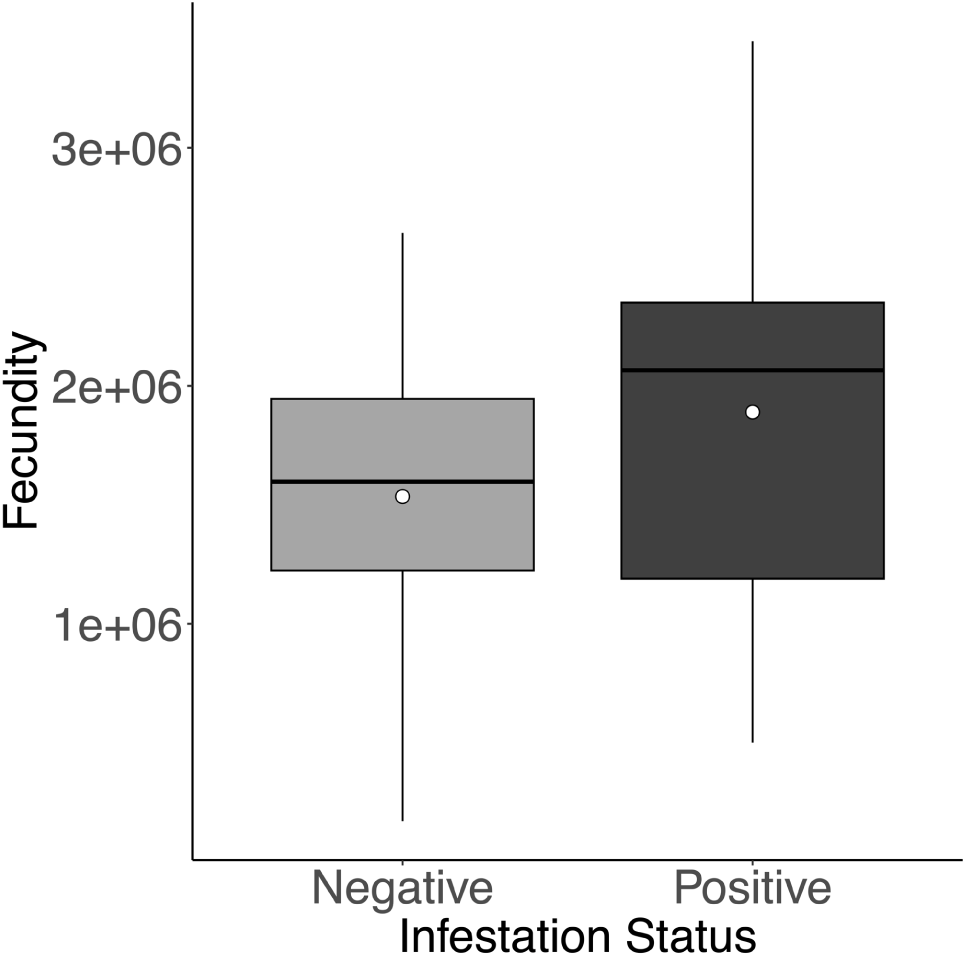
Effect of infestation status on fecundity. General linearized model, p-value = 0.577. The middle line of the box plot represents the median, and the white dot represents the mean.

**Table 1.** Parameter estimates for the predictors in the generalized linear model of fecundity.

| Parameter | Estimate +/- SE | *t* value | *P* value |
| --- | --- | --- | --- |
| Intercept | -2.81e+05 ± 3.69e+05 | -0.762 | 0.45 |
| Crab Body Mass | 6.15e+04 ± 1.09e+04 | 5.634 | <0.0001 |
| Carapace Width | 2.78e+04 ± 7.19e+03 | 3.867 | 0.0003 |
| Embryo Stage | 1.11e+05 ± 2.55e+05 | 0.434 | 0.666 |
| Infestation Status | 3.26e+05 ± 5.80e+05 | 0.562 | 0.577 |
| Embryo Stage x Infestation Status | -9.05e+04 ± 3.50e+05 | -0.259 | 0.797 |

The mean reproductive output for all locations was 0.28 ± 0.08 g of viable embryo mass per gram of body mass, not including legs. The most significant predictor of reproductive output was egg stage (Fig. 4, 5, Table 2). Crabs with early-stage embryos had a higher reproductive output than crabs with late-stage embryos. Infestation status was not a significant predictor of reproductive output (Fig. 5, Table 2).

**Figure 4.**
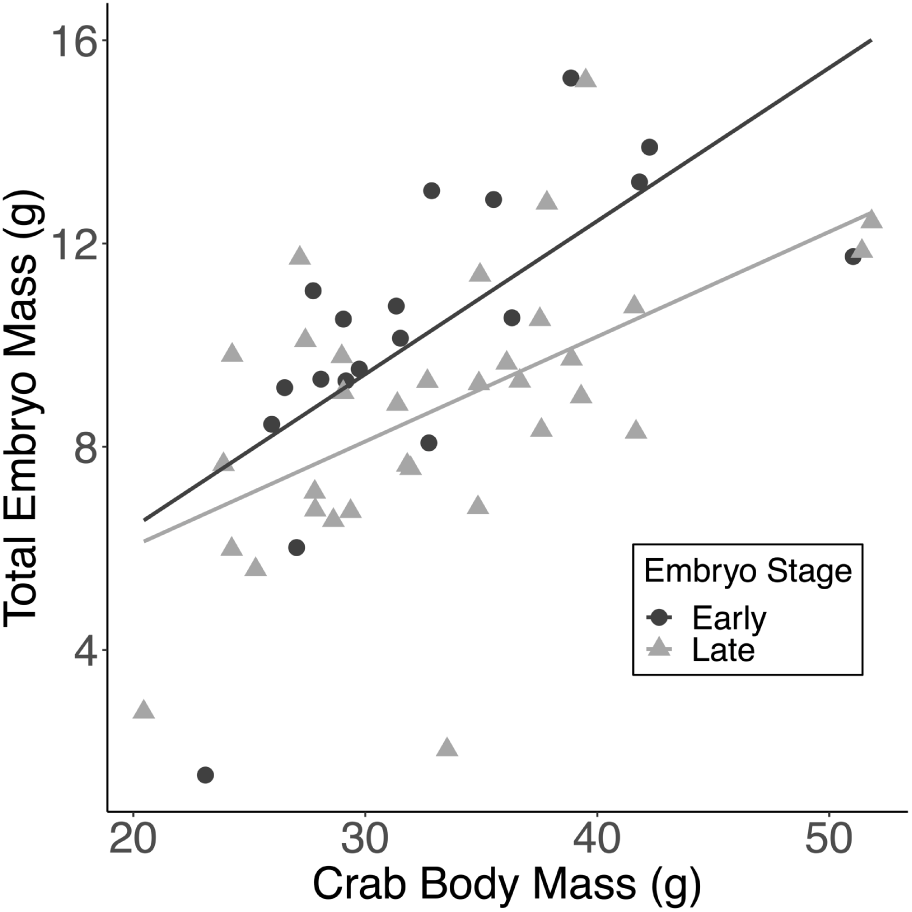
Effect of embryo stage on the relationship between embryo mass and total body mass, or reproductive output. General linearized model, p-value <0.05. The slope, or total embryo mass divided by body mass, represents reproductive output, and is greater for crabs with early-stage embryos.

**Figure 5.**
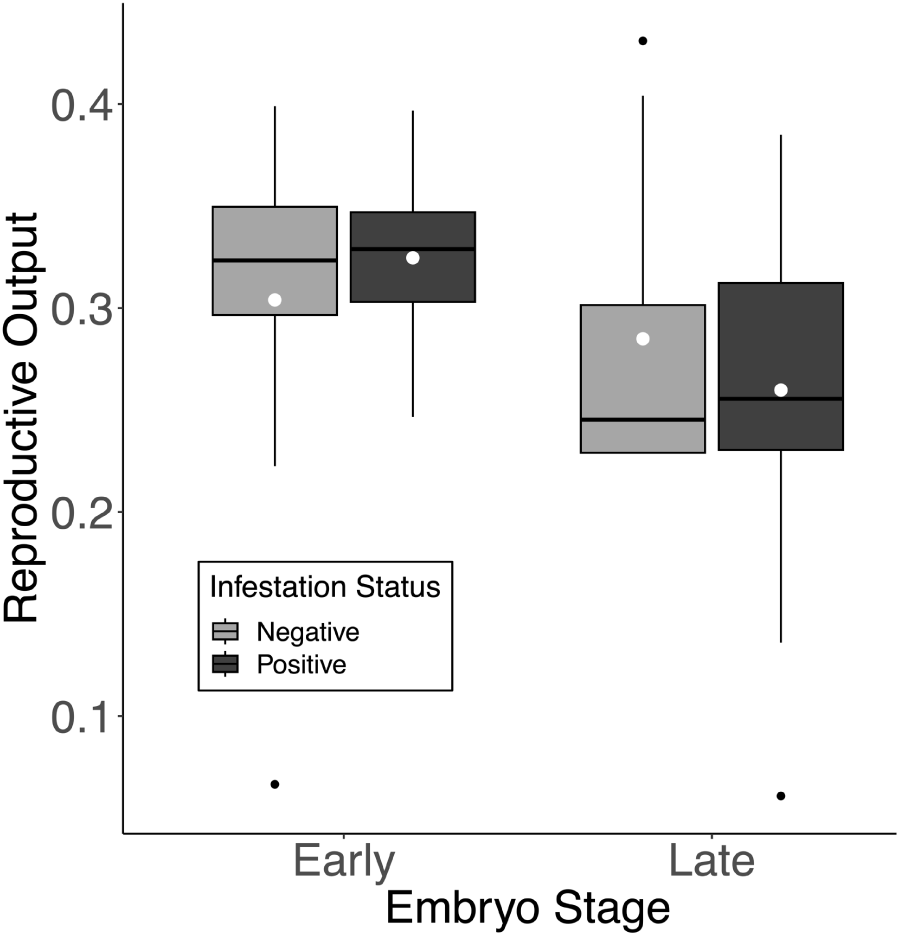
Effect of infestation status and embryo stage on reproductive output. General linearized model, p-value >0.05 (infestation status), p-value <0.05 (embryo stage). The middle line of the box plot represents the median, and the white dot represents the mean. Black dots represent outliers.

**Table 2.** Parameter estimates for predictors in the generalized linear model of reproductive output.

| Parameter | Estimate +/- SE | *t* value | *P* value |
| --- | --- | --- | --- |
| Intercept | 0.35726 +/- 0.03681 | 9.705 | <0.0001 |
| Embryo Stage | -0.04569 +/- 0.0216 | -2.115 | 0.039 |
| Infestation Status | 0.0732 +/- 0.08183 | 0.895 | 0.376 |
| Embryo Stage x Infestation Status | -0.048337 +/- 0.04846 | -0.997 | 0.324 |
| Nemertean Density | -0.0693 +/- 0.31973 | -0.217 | 0.829 |

Embryo stage was a significant predictor of infestation status (Fig. 6). Out of the 19 crabs with early-stage embryos, only 7 were infested with nemerteans. Out of the 33 crabs with late-stage embryos, 25 were infested with nemerteans. Average nemertean density in worms with positive signs of infestation was 0.72 ± 1.39 worms per 1000 embryos. Out of the 32 worms with positive signs of infestation, 12 had a nemertean density of 0 worms per 1000 embryos because no worms were found, only mucus tubes or eaten embryos. Infestation status was a significant predictor of nemertean density (Table 3). Embryo stage was not a significant predictor of nemertean density (Fig. 7, Table 3).

**Figure 6.**
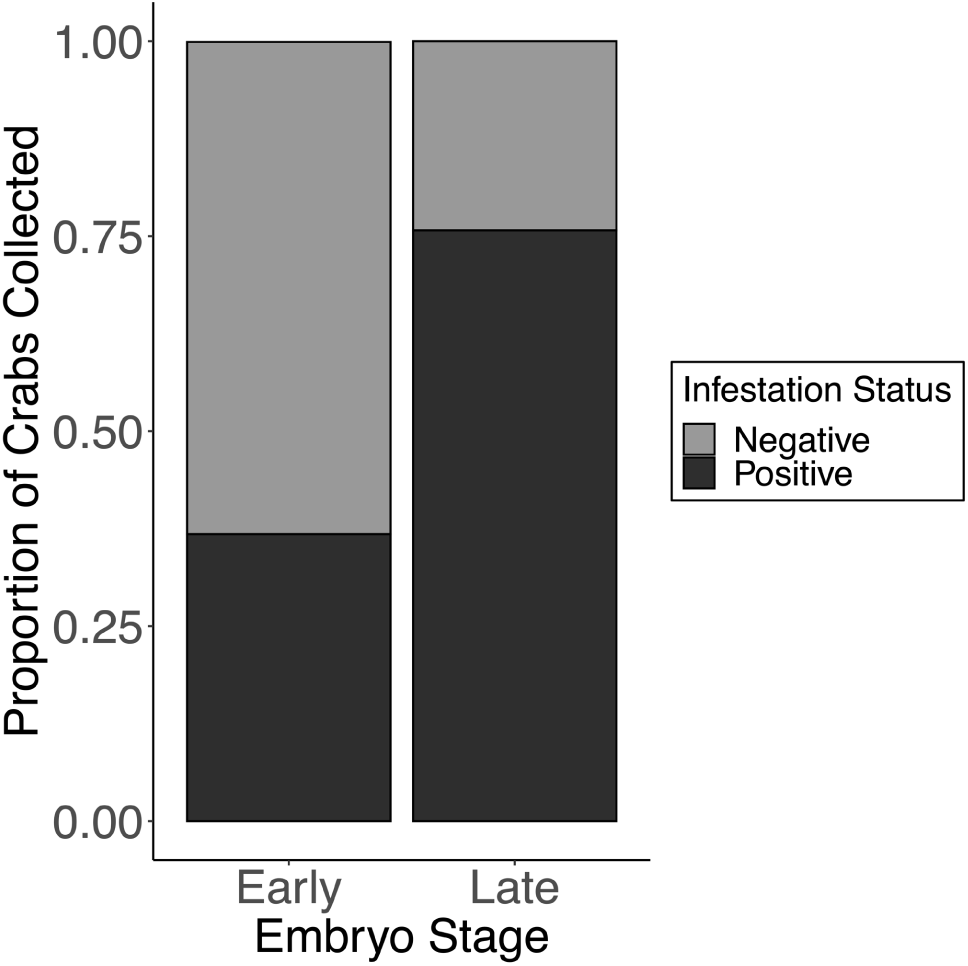
Proportion of crabs collected with *C. carcinophila* infestations as affected by crab embryo stage.

**Figure 7.**
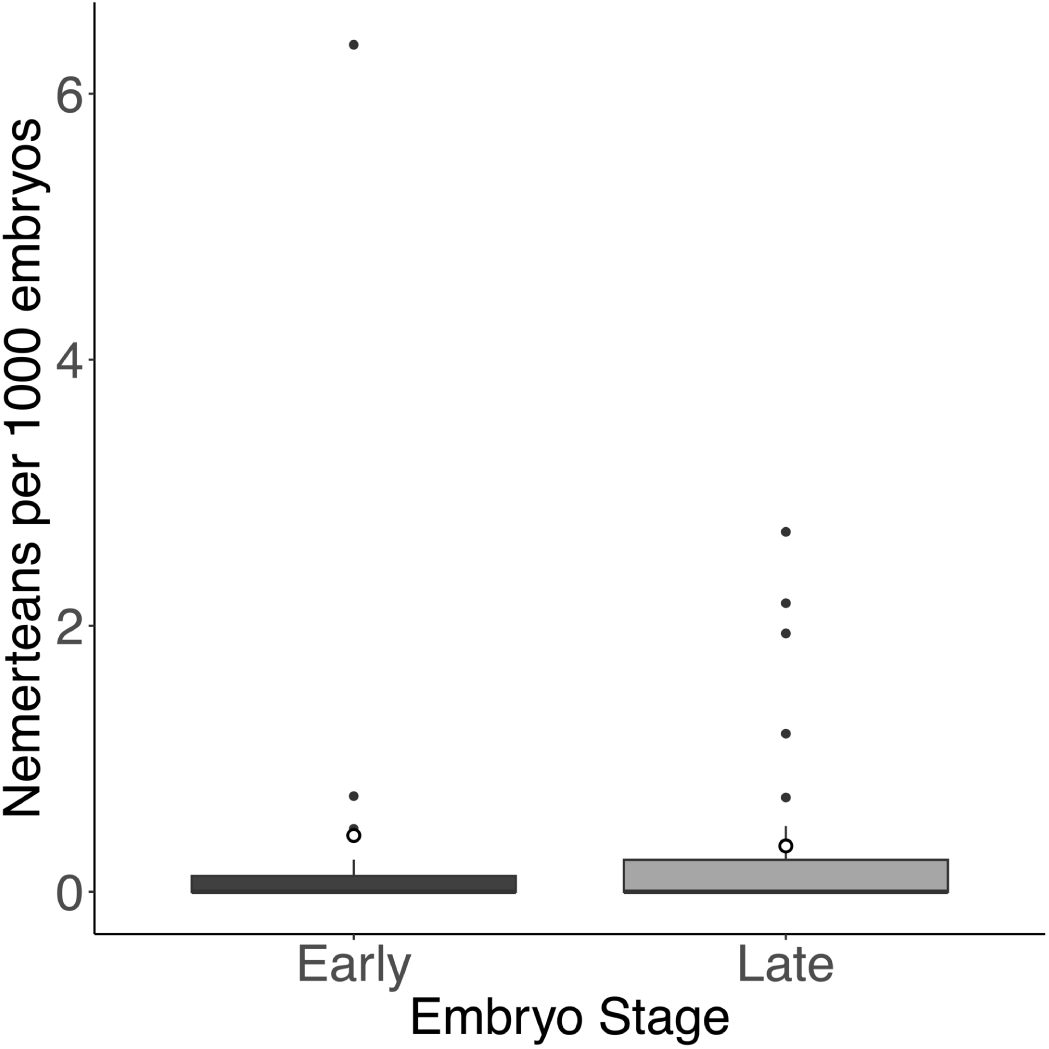
Effect of embryo stage on nemertean density. General linearized model, p-value = 0.557. The middle line of the box plot represents the median, and the white dot represents the mean. Black dots represent outliers.

**Table 3.**
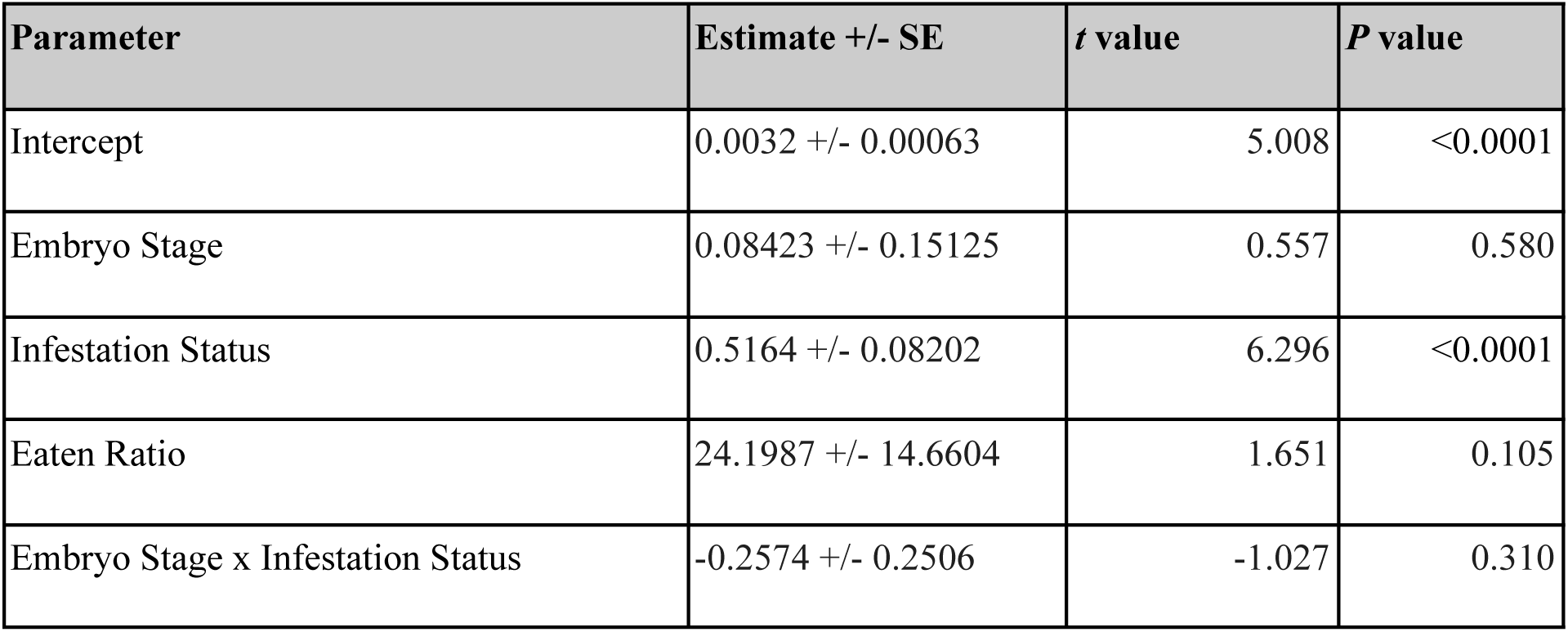
Parameter estimates for predictors in the generalized linear model of nemertean density (worms found per 1000 embryos).

| Parameter | Estimate +/- SE | *t* value | *P* value |
| --- | --- | --- | --- |
| Intercept | 0.0032 +/- 0.00063 | 5.008 | <0.0001 |
| Embryo Stage | 0.08423 +/- 0.15125 | 0.557 | 0.580 |
| Infestation Status | 0.5164 +/- 0.08202 | 6.296 | <0.0001 |
| Eaten Ratio | 24.1987 +/- 14.6604 | 1.651 | 0.105 |
| Embryo Stage x Infestation Status | -0.2574 +/- 0.2506 | -1.027 | 0.310 |

Average number of eggs eaten per 1000 live embryos was 28.44 ± 52.36 eggs. Infestation status and nemertean density were significant predictors of the ratio of eaten eggs to live embryos (Fig. 8, Table 4). Embryo stage was not a significant predictor of the ratio of eaten eggs to live embryos (Table 4).

**Figure 8.**
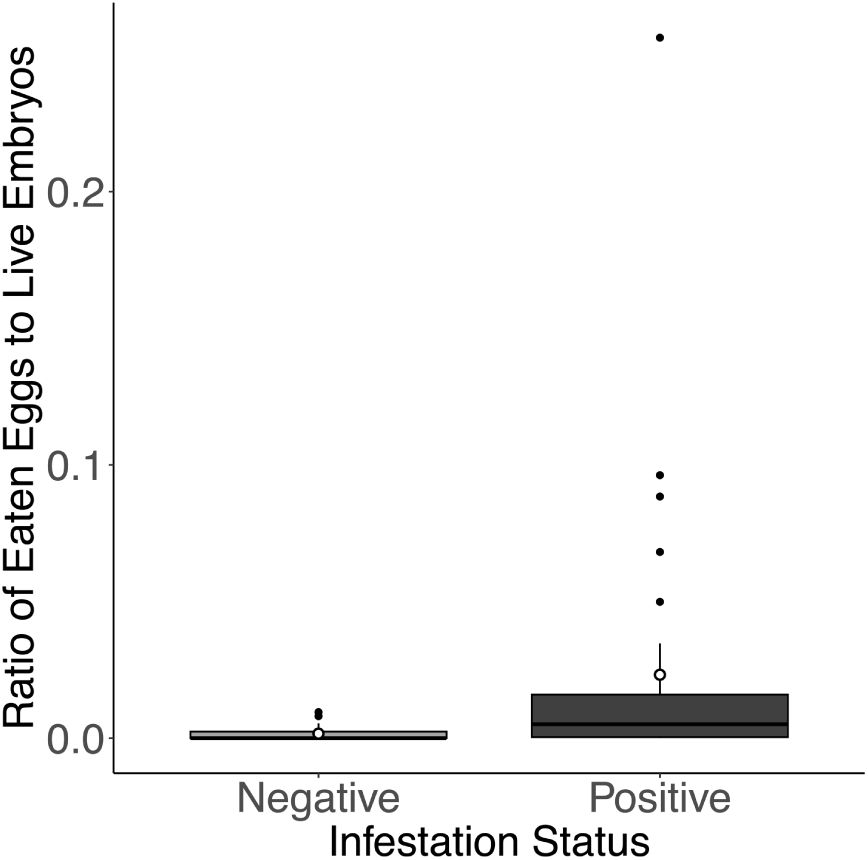
Effect of infestation status on ratio of eaten eggs to live embryos. General linearized model, p-value <0.05. The middle line of the box plot represents the median, and the white dot represents the mean. Black dots represent outliers.

**Table 4.** Parameter estimates for predictors in the generalized linear model of the ratio of eaten eggs to live embryos.

| Parameter | Estimate +/- SE | *t* value | *P* value |
| --- | --- | --- | --- |
| Intercept | -4.3728 +/- 0.3054 | -14.318 | <0.0001 |
| Egg Stage | 0.3114 +/- 0.285 | 1.093 | 0.275 |
| Infestation Status | 0.6288 +/- 0.283 | 2.222 | 0.026 |
| Nemertean Density | 0.5224 +/- 0.2518 | 2.074 | 0.038 |
| Egg Stage x Infestation Status | 0.429 +/- 0.61774 | 0.695 | 0.487 |

## DISCUSSION

With new nemertean species discovered each year, it becomes increasingly important to assess the ecological and economic impacts of those already known. Our study addresses this need by quantifying the effect of a single species, *Carcinonemertes carcinophila*, on the reproductive capacity of its host, *Callinectes sapidus*. While we found that infestation status did not significantly affect overall fecundity or total reproductive output, other biological variables played key roles. Carapace length and crab body mass were both significant predictors of fecundity, while embryo stage did not show a significant relationship. However, reproductive output was strongly influenced by embryo stage, with crabs carrying early-stage embryos exhibiting higher outputs. Interestingly, nemertean infestations were more common in crabs bearing late-stage embryos. Both nemertean density and infestation status were significant predictors of the ratio of eaten eggs to live embryos, suggesting that while total fecundity may remain stable, the presence of nemerteans can compromise embryo survival. These findings highlight the need for more comprehensive assessments of nemertean impacts across crustacean hosts and ecosystems.

To place our findings in a broader geographical context, we compared the prevalence of *C. carcinophila* in Florida with previous reports from Chesapeake Bay. The prevalence of *C. carcinophila* in Florida was lower than the prevalence in Chesapeake Bay measured by

Schneider et al. (2023). This may be because Schneider et al. 2023 accounted for worms infested in the gills. Around 7.5% of infested crabs in the Chesapeake Bay study found worms solely in the gills. This study dealt with the impact of worms infested in the brood mass on fertility. Gill infestations were not included due to their logistical difficulty.

Beyond regional differences in prevalence, we also observed a clear temporal pattern in infestation dynamics. In our study, we observed the rate of infestation to increase over the duration of the embryonic period, the same pattern that most nemertean egg predators follow (Kuris and Wickham 1987, Shields 1993, Kuris 1978). Crabs in late embryogenesis tested positive for infestation more than those in early embryogenesis. This is most likely due to the nemertean life cycle; when a female oviposits her eggs, the worms start making their way from the gills to the brood mass. Later in embryogenesis, the worms have had more time to travel to the brood mass, lay eggs and eat embryos, which all led to a positive determination of infestation. Although worms have an easier time consuming recently oviposited eggs (Okazaki 1986, Roe 1984), temporal factors play a greater role in controlling patterns of egg consumption (Kuris and Wickham 1987).

There are a few key differences between the *C. carcinophila* blue crab relationship and other decapod species that have been more severely impacted by nemertean infestation (Table 5), such as Dungeness or red king crabs. The first difference is in the infestation intensity, where similar estimations of worms per brood mass exist between blue crabs and Dungeness or red king crabs, but the number of worms per embryo was much lower for *C. sapidus* in Florida. At the peak of *C. errans* infestations in the late 1970s, the nemertean intensity ranged from 8.56 to 24.96 worms per 1000 embryos (Wickham et al. 1979). In red king crabs, *C. regicides* were reported at intensities greater than 10⁴ per pleopod (Kuris et al. 1991). While red king crabs exhibit considerably higher infection intensity, both host species had much higher nemertean densities than *C. carcinophila* in blue crabs.

Another key difference between C. *carcinophila* and other systems lies in the host embryonic period, defined as the number of days between egg extrusion and hatching. The embryonic period of the Florida blue crab is relatively short, lasting around 12 d in temperatures 28°C and greater, but has been studied to take longer, around 14-18 d, at colder temperatures (Darsono 1992, Costlow and Bookhout 1959). The embryonic period of the red king crab is variable based on parity, where primiparous crabs’ embryonic period lasts around 365 days and multiparous crabs last for 328 days (Stevens et al. 2007). Dungeness crabs also have a much longer embryonic period than the blue crab, around 65 to 130 days (Rasmuson 2013). The considerably longer embryonic periods of Dungeness and red king crabs suggest more time for nemerteans to prey on brood masses, resulting in more eggs consumed and a greater percentage of the crabs’ fecundity lost to egg predators.

Egg consumption rates also differ between *C. carcinophila* and other nemertean egg predators. *C. carcinophila* is estimated to consume 0.16-4.5 eggs per day throughout the blue crab’s 12-day embryonic period, or about 0-0.0044% of the brood mass (Schneider et al. 2023). While *C. errans* consumes a comparable 0.5-1.5 eggs per day (Hamilton, 1984), their impact is exacerbated by the Dungeness crab’s 90-day embryonic period and has been linked to 55% of the brood mass loss in *C. magister* (Wickham 1979). In the yellow shore crab *Hemigrapsus oregonensis,* the nemertean *Carcinonemertes epialti* consumes on average 1.02-1.67 eggs per day and causes 7.5-19% of brood loss (Kuris and Wickham 1987). In spiny lobsters, *C. conanobrieni* consumes up to 43.8% of brood masses (Berben et al. 2023).

A final key difference between these crab-nemertean interactions is not the feeding rates, but the density of worms found in each crab. Higher worm densities, though mostly influenced by the host embryonic period and total fecundity, could also suggest different reproductive strategies. Nemertean reproductive strategies seem to be divided on the basis of significant brood mortality. Parthenogenesis, or the ability to produce offspring without fertilization from a male, is present in *C. errans* and *C. epialti* (Roe, 1986), both of which can cause significant brood mortality in *C. magister* and *H. oregonensis,* respectively. Meanwhile, *C. carcinophila* is only able to produce eggs via external fertilization (Wickham and Kuris 1985). A more difficult reproductive process may result in less explosive worm populations. All of this is impacted by the host’s embryogenic period. In *C. sapidus,* that period is too short for the slow feeding rate of *C. carcinophila* to impact its fertility.

Nemertean density estimates were much higher than those in the Chesapeake Bay. The estimated nemerteans per brood mass was 735, significantly higher than the Chesapeake Bay estimates of 46-62 worms per brood mass. The observed outliers in nemertean density suggests an aggregated pattern typical of parasites, consistent with a negative binomial distribution in which most hosts experience low infestation, but a minority bear exceptionally high burdens. The higher nemertean density in Florida could be due to differences in reproduction between Chesapeake Bay and Florida nemerteans or their blue crab hosts. Florida blue crabs are reproductive year-round, unlike Chesapeake Bay crabs that undergo an overwintering period. Warmer water allows Floridian blue crabs to produce more broods, which could possibly increase nemerteans. Warmer temperatures can also affect nemertean feeding rates and oviposition (Shields and Kuris 1988). A study on *Carcinonemertes errans,* the nemertean that preys on the Dungeness crab, is affected by salinity (Dunn et. al 2015). Low-salinity treatment decreased survival rates of *C. errans* larvae. Though secondary to salinity, temperature was also found to have a significant impact on larval survival (Dunn 2011). Other studies have linked higher temperatures to increased nemertean larval survival at low salinities (Zhao and Sun 2006, Charmantier et. al 1991). In some areas, Floridian estuaries are recorded to have higher salinities than those in the Chesapeake, due to the Chesapeake’s higher freshwater inputs (Xu et al. 2012) and Florida’s greater tidal influence (Guenther 2012). However, it should be noted that almost no studies compare the salinities of Floridian and Chesapeake estuaries, and that existing studies note the highly variable nature of estuarine environments. Florida’s higher salinities and temperatures could have been favorable to the *C. carcinophila* larvae, resulting in higher survival of parasite larvae, further maturation of worms and ultimately higher observed nemertean density in the brood mass. More experimentation on salinity and temperature stressors should be done on *C. carcinophila* to determine if this is true.

Other aspects of *C. carcinophila* reproduction are also deserving of future studies. There have been no fecundity estimates for *C. carcinophila* as it relates to the embryonic period of *C. sapidus.* Parasite intensity is directly related to loss of host fecundity (Kuris and Wickham 1987). A higher parasite fecundity could help explain why species like *H. oregonensis* have higher densities of infestation, and therefore greater brood mortality than *C. sapidus* (Table 5). Future studies should also aim to replicate the parthenogenesis experiments of Roe (1986) in multiple species from the *Carcinonemertes* genus. Ease of reproduction allowed by parthenogenesis could also contribute to higher parasite intensities and greater brood mortality.

## CONCLUSIONS

Similar to results discussed in Schneider et al. 2023, *C. carcinophila* appeared to have little impact on the reproductive output of Florida blue crabs. However, the geographic and environmental scope of our study was limited and several findings warrant further investigation, particularly the much higher infestation intensity among Florida blue crabs. Perhaps there are other areas of the Gulf of Mexico or the Atlantic Coast where this high infestation intensity is associated with impacts to reproduction. Future research efforts should also be turned to discovering those that might pose a threat to commercially important crustacean fisheries. The Dungeness and red king crabs have shown us that nemertean worms can pose an economic and ecological threat to decapod crustaceans. Exploring the fertility impact of emerging parasitic species could help mitigate that potential threat. Impacts of newly described nemertean species, such as *Carcinonemertes camanchaco,* are still unknown and merit considerable research effort. From Maine to Argentina, commercial crustacean fisheries are vulnerable to the danger of egg predators.

## ACKNOWLEDGEMENTS

We thank Todd Hoy and Raymond King for providing the blue crabs for this study. The University of Florida College of Liberal Arts and Sciences’ University Scholars Program provided the funding to conduct this study. We also thank Ryan Bergman, Chris Skibicki, Brynn Fricke, Emma Kavanaugh and Sam K for their help processing crabs and performing subsample counts.

